# UniFuncNet: a flexible network annotation framework

**DOI:** 10.1101/2022.03.15.484380

**Authors:** Pedro Queirós, Oskar Hickl, Susana Martínez Arbas, Paul Wilmes, Patrick May

**Affiliations:** Systems Ecology, Luxembourg Centre for Systems Biomedicine, University of Luxembourg, 4367, Luxembourg; Bioinformatics Core, Luxembourg Centre for Systems Biomedicine, University of Luxembourg, 4367, Luxembourg; Department of Life Sciences and Medicine, Faculty of Science, Technology and Medicine, University of Luxembourg, Esch-sur-Alzette, Luxembourg

## Abstract

**Summary:** Functional annotation is an integral part in the analysis of organisms, as well as of multi-species communities. A common way to integrate such information is using biological networks. However, current data integration network tools are heavily dependent on a single source of information, which might strongly limit the amount of relevant data contained within the network. Here we present UniFuncNet, a network annotation framework that dynamically integrates data from multiple biological databases, thereby enabling data collection from various sources based on user preference. This results in a flexible and comprehensive data retrieval framework for network based analyses of omics data. Importantly, UniFuncNet’s data integration methodology allows for the output of a non-redundant composite network and associated metadata. In addition, a workflow exporting UniFuncNet’s output to the graph database management system Neo4j was implemented, which allows for efficient querying and analysis.

**Availability:** Source code is available at https://github.com/PedroMTQ/UniFuncNet.

## Introduction

There exists an unprecedented amount of biomolecular data available thanks to the advances in, among others, sequencing, mass spectrometry and bioinformatics techniques. This allows for the study of function across several biological levels at high resolution, from single organisms to the combined functional potential of microbial communities. This wealth of information is difficult to access and use in a straightforward and scalable manner, e.g., due to the lack of a universal data repository and the use of a multitude of data formats and annotations.

Networks are frequently used for large-scale omics data analyses as these are versatile tools that can be used to model complex biological systems [1]. The identification and mapping of functional entities (e.g., proteins) to networks are central tasks performed during large-scale studies of new species or microbial communities. For example, networks have been used to study ecological interactions such as metabolic cross-feeding, synergism, and antagonism [2], to detect correlations in metabolic networks [3], to identify keystone functions and genes [4].

Given the available functional annotations linked to omics data, a common modelling approach, among others [5], is to use genome-scale metabolic models (GSMM) to integrate all, or part, of the metabolic and transport reaction network(s) within an organism or community [6, 7]. Such networks are usually derived by mapping functional annotations to the corresponding reactions and pathways [8, 9], and can be used for the *in silico* simulation of metabolism.

Several methodologies [10] and tools [11] are now available for the automated generation and semi-curation of GSMMs. Many methods are able to automatically and accurately reconstruct well-known parts of metabolism, which, due being shared by many taxa [12], have been more extensively studied [13]. While the apparent conservation in function based on homology is advantageous when modelling well-studied metabolism, the resulting GSMMs are often very general and redundant, which may not capture the peculiarities of individual organisms. Modelling species-specific metabolic pathways is important, e.g., for understanding microbial interactions [14], but challenging, since annotations are often incomplete [15, 16]. Here, knowledge integration from multiple databases (e.g., MIBig [17], KEGG [18], and MetaCyc [19]) may help.

Even though some resources provide frameworks for mapping functional entities (e.g., KEGG [18] and MetaCyc [19]), combining them into a more comprehensive resource at a case-by-case basis is laborious, since this integration requires extensive cross-linking, and often manual review/curation. One additional complication is the use of different ontologies [20], which leads to the necessity of cross-linking ontology systems with varying structures and resolutions (e.g., KEGG orthologs and gene ontologies[21, 22]). Additionally, while some databases are structured and provide access through the use of application programming interfaces (API) (e.g., KEGG), relational or non-relational or other standardized formats (e.g., json and xml), others provide data in semi-unstructured formats, thereby requiring the implementation of more specialized data processing methodologies (e.g., text mining [23]).

In essence, the diversity and quantity of biological databases, constitute some of the major challenges in the integration of such data. These, and other technical challenges, make such resources inaccessible to researchers without a computational background.

The challenge of integrating knowledge from multiple sources in an automated manner in the context of network analysis was tackled through the development of the presented network annotation framework - (Uni)fied (Func)tional (Net)work (UniFuncNet). UniFuncNet automates the highly time-consuming process of searching multiple databases, extracting and integrating data into a composite output. Biological databases commonly contain multiple entry types (e.g., compounds or genes), therefore, UniFuncNet’s implementation reflects the general structure of such databases; for this purpose, we modelled four different entity types: genes, reactions, proteins, and compounds. In turn, these data models can then be linked as a network, and used for storing and exporting information in machine and human-readable formats. Combining data models with multiple data collection methodologies results in a flexible yet robust data retrieval framework. In turn, this allows researchers to fine-tune UniFuncNet to their specific routine data integration tasks, starting from simple use cases such as collecting ChEBI identifiers (IDs) for a list of compound names and finding reactions for certain protein IDs to linking compounds to organisms, or expanding GSMMs. To showcase how the user can include UniFuncNet in their analysis, the last two previously mentioned use cases have been implemented as separate example workflows; while these are simple wrappers around UniFuncNet and other tools, they may serve as a template for future, and potentially more complex, workflows.

UniFuncNet aims to provide a straightforward, versatile, and accessible data collection and network annotation framework. UniFuncNet will prove useful across multiple domains of bioinformatics, especially at a moment in time where large-scale data integration is seen as fundamental rather than optional. In order to provide an easily and efficiently queryable database, we implemented an API that exports UniFuncNet’s data to Neo4j.

## Materials and methods

### Implementation

UniFuncNet was implemented in Python (v3.9) and currently collects data from KEGG [18], MetaCyc [19], Rhea [24], ChEBI [25], HMDB [26], UniProt [27] and Pubchem [28], cross-linking the information between these databases. For web data collection, UniFuncNet uses the Python package “requests” (v2.25.1), which queries each database and collects the respective response (usually HTML or json). To parse the HTML responses the “beautiful soup” [29] package is used (v4.10.0). For some of the databases, i.e., MetaCyc [19], Rhea [24] and ChEBI [25] the database flat files are first downloaded, parsed and stored locally in a SQLite (v3.36.0) database. In order to use the MetaCyc database, the user must obtain a license (academic licenses are freely available) from MetaCyc (which we recommend since it’s a highly curated and comprehensive resource). For web data collection, UniFuncNet makes use of API calls to retrieve information (if possible). However, whenever necessary, data is collected by querying the database’s website and parsing the query result (i.e., web scraping). Each query result (web or local data) is parsed according to the step of the workflow and database being queried (with database-specific scrapers), and standardized according to UniFuncNet’s framework. This data parsing allows for the retrieval of annotations (IDs and synonyms) as well as any connections between database entries.

To avoid overloading the respective web servers, UniFuncNet works in a strictly sequential manner and additionally enforces a time-out for requests to the same server (10 seconds in-between queries by default). Additionally, in order to avoid repeating web queries, UniFuncNet saves past web queries in memory and retrieves the necessary entity whenever a query is repeated. This sequential methodology has the additional benefit of not creating redundant entities which may lead to downstream issues with redundancy and output network connectivity.

The results shown in this paper were collected from multiple sources, MetaCyc version 25.1 was used; the Rhea and ChEBI data corresponded to the flat files uploaded on the 17th of November, 2021; and all data extracted from the multiple websites was collected on the 24th of January, 2022. The version of UniFuncNet used in this paper is v1.02.

### Input and output

UniFuncNet takes as input a tab-separated file, containing a list of IDs (e.g., “P02769”), ID types (source of the IDs, e.g., “uniprot”), entity types (e.g., “protein”), and search modes (e.g., “pg”, for “protein-to-gene”).

UniFuncNet outputs one tab-separated file per entity type, i.e., genes, proteins, reactions, and compounds, listing all the searched entities along with any associated metadata (e.g., database IDs) and all the associations between each entity. Additionally, it outputs a file in simple interaction format (SIF), which allows for integration into network frameworks, such as Cytoscape [30] or Neo4j.

For a detailed description of input format requirements and all outputs, as well as a usage guide refer to UniFuncNet’s documentation at https://github.com/PedroMTQ/UniFuncNet.

### Workflows methodology

We implemented two example workflows to showcase potential applications of UniFuncNet. The first workflow relates to the expansion of GSMMs using UniFuncNet, and the second to the mapping of compounds to organisms. An example use case is provided for each of these workflows. In these use cases, UniFuncNet collected information from the databases KEGG, MetaCyc, Rhea, and ChEBI. The code used for the generation of results is available at https://gitlab.lcsb.uni.lu/pedro.queiros/benchmark_unifuncnet. After installation and download of the required tools and data, the workflows are fully automated (e.g., automatically launching tools and doing the necessary data processing). Mantis [31] v1.3 was run for the functional annotation, using the KOfam [32], Pfam [33] and MetaCyc [19] reference databases (the MetaCyc database was generated with the code in https://github.com/PedroMTQ/refdb_generator).

### Workflow I - Expansion of GSMMs

This workflow (Figure 1.A) receives as input multiple protein fasta files, i.e., proteomes, and outputs an expanded network per sample in SIF format. The proteomes are passed to Mantis [31] while GSMMs are created with CarveMe [34]. The enzyme commission numbers (ECs) and MetaCyc protein IDs absent in the CarveMe GSMMs are exported from the Mantis’ functional annotations into a UniFuncNet-formatted input file (using the “prc” search mode). In this manner UniFuncNet can be used to collect data on the additional ECs and MetaCyc protein IDs and connect them to the original GSMMs. In order to exclude unspecific interactions, edges connecting to common cofactors were removed (this list of cofactors has been manually curated but can be edited and is available at https://github.com/PedroMTQ/UniFuncNet/tree/main/Resources/cpd_to_ignore.tsv).

**Figure 1.**
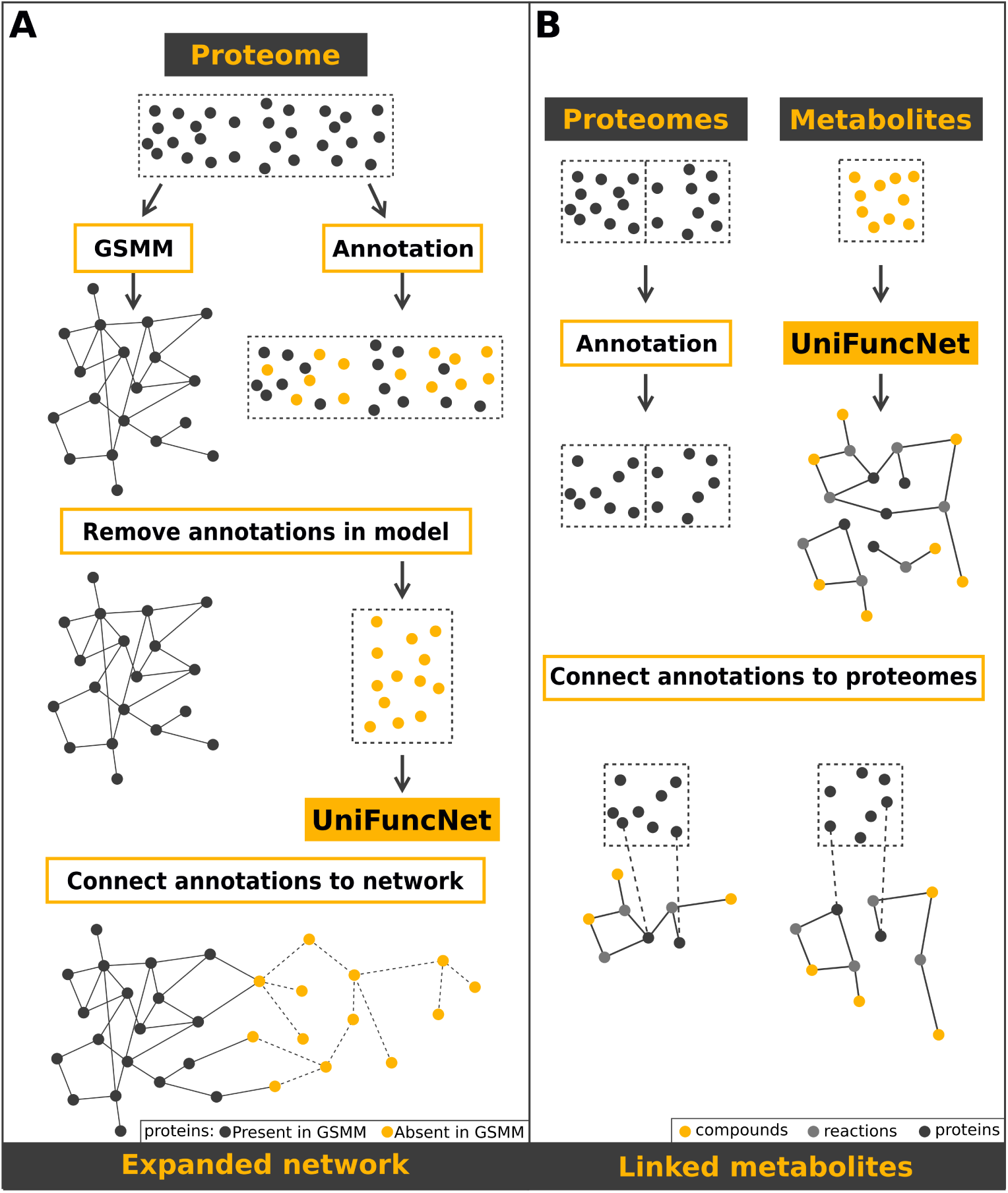
Use cases workflows. **A** Workflow I: UniFuncNet is used to aid in the expansion of a previously generated GSMM. First, a draft GSMM (grey network) and functional annotations (dashed box with grey and yellow nodes) are extracted from the input proteome (top dashed box with grey nodes). Next, all functional annotations absent (dashed box with yellow nodes) in the model are input into UniFuncNet. Lastly, all of the metabolic model’s entities are connected to UniFuncNet’s output (yellow and grey nodes connected with non-dashed edges). Optionally, the user may also add all remaining nodes in UniFuncNet’s network (yellow nodes connected with dashed edges). **B** Workflow II: UniFuncNet is used to identify the proteins of an organism involved in the metabolism of specific compounds. First, proteomes for all organisms were collected (represented by the first dashed box with black dots), these were then functionally annotated with Mantis (represented by the second dashed box with black dots, note the lower number of nodes in each proteome, which represents the lack of functional annotations for some proteins). Using UniFuncNet, we created a network with the reactions and respective proteins associated with each input compound. Lastly, using the previously created network, we linked the compounds with their respective proteins in each proteome.

The workflow was implemented with CarveMe [34] v1.5. Additional information is available at https://github.com/PedroMTQ/UniFuncNet/tree/main/Workflows/GSMM_Expansion. It is crucial to note that this workflow is merely an example use case, therefore any output files created should be thoroughly curated.

As an example application (henceforth referred to as “use case I”), we used the following five organisms’ SwissProt[27] reference proteomes: UP000001031 [35] for *Akkermansia muciniphila*, UP000025221 [36] for *Bradyrhizobium japonicum*, UP000018291 [37] for *Microthrix parvicella*, UP000002528 [38] for *Pelagibacter ubique*, and UP000000586 [39] for *Streptococcus pneumoniae*.

To evaluate the functional redundancy of the baseline and expanded networks, a presence/absence encoding of each network’s ECs was applied, followed by a cosine distance calculation using the NLTK package (v3.5), where equal encoded vectors have a score of 1 and completely different a score of 0. This calculation is henceforth referred to as the “ECs functional redundancy”.

### Workflow II - Omics cross-linking

The second workflow (Figure 1.B) attempts to link compounds to specific organisms by searching for information on compounds and linking them to functionally annotated organisms. The input are multiple species proteomes and information (i.e., IDs) on the compounds of interest. The output is a network connecting each compound to all proteomes, and hence, to all organisms. Additional information is available at https://github.com/PedroMTQ/UniFuncNet/tree/main/Workflows/Compounds_to_Organisms_Mapping.

As an example application of this workflow (henceforth referred to as “use case II”), we applied it to the metabolomics dataset MTBLS497 from Metabolights [40]. In the respective experimental study [41] four organisms, *Escherichia coli, Klebsiella pneumoniae, Pseudomonas aeruginosa*, and *Staphylococcus aureus*, were cultured, sampled and analysed to link them with 13 compounds of interest. The following proteomes from UniProt were used: *E. coli* UP000001410 [42], *K. pneumoniae* UP000000265 [43], *P. aeruginosa* the proteome UP000002438 [44], and *S. aureus* UP000008816 [45].

## Results

### UniFuncNet

UniFuncNet is a network annotation framework that collects user-defined data from multiple biological databases, e.g., KEGG orthology IDs (Figure 2). The user input determines which information is collected by UniFuncNet. UniFuncNet retrieves data from the respective biological databases and parses it; if applicable, it then branches out and gathers any additional data associated with the originally retrieved data. This is repeated iteratively until all sources of information are exhausted. When retrieving information for compounds, UniFuncNet can retrieve data based on synonyms, and not only IDs, as it may facilitate the integration of data where only synonyms are available. However, the reliability of synonyms-based data retrieval is inferior to IDs due to its ambiguity [46]. UniFuncNet is available as a conda package, and it’s respective documentation is available at https://github.com/PedroMTQ/UniFuncNet.

**Figure 2.**
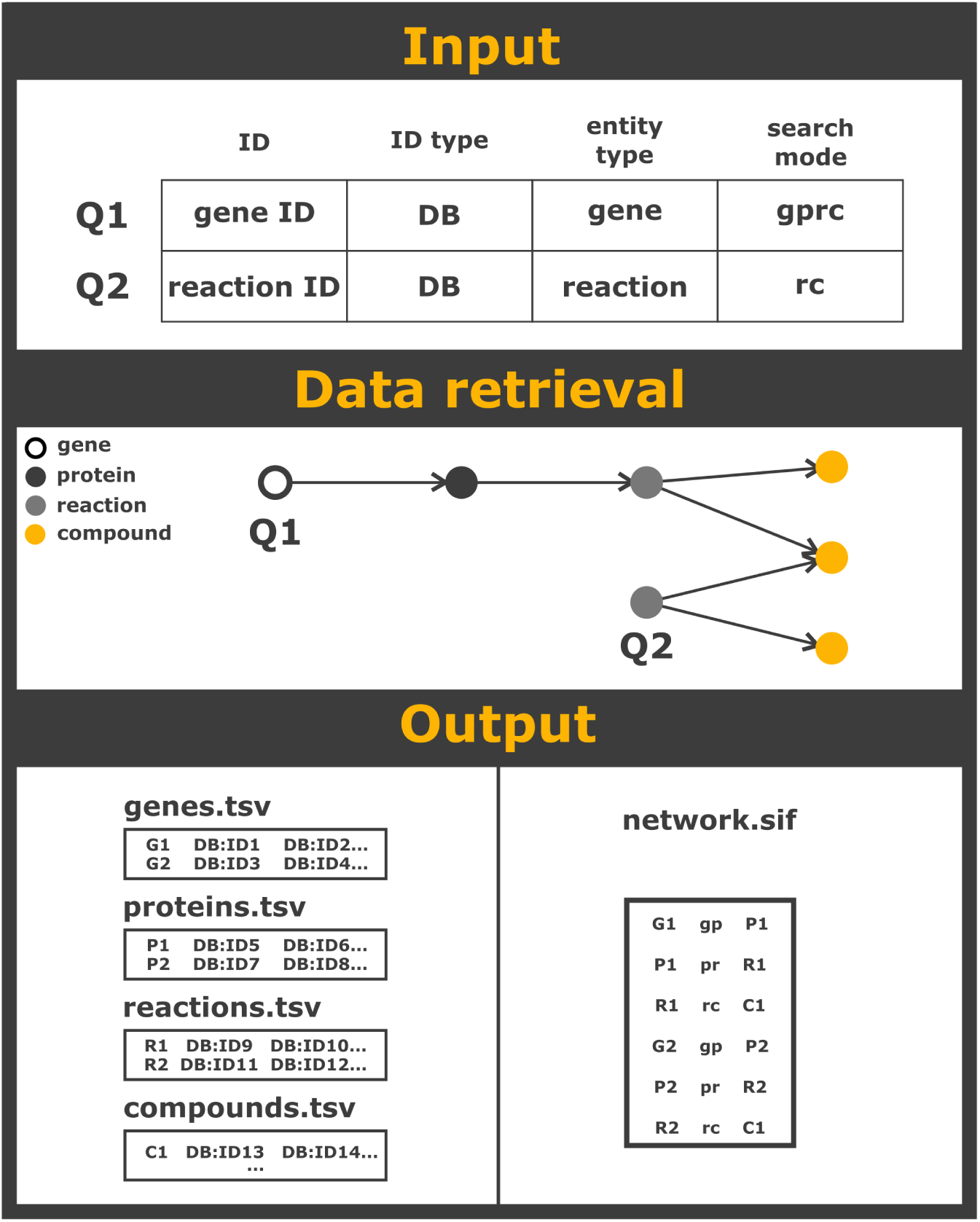
UniFuncNet overview. The input of UniFuncNet is a list of IDs, ID types, entity types, and search modes, which are processed line by line. In this example, UniFuncNet starts by collecting data on the first query (Q1), which is a gene. According to the search mode “gprc” it then searches for data for the connected proteins, reactions and compounds. For the second query (Q2) - a reaction, UniFuncNet first collects data on the reaction and then on the associated compounds (search mode “rc”). UniFuncNet then outputs the results for each collected entity in the respective tsv file, as well as the resulting network in SIF format.

#### Data models

In order to standardize the representation of the multiple types of data within the UniFuncNet framework, we implemented multiple data models, each one representing an entity type, i.e., compounds, reactions, proteins, and genes. In general, entities are associated with IDs from multiple databases and other entity-specific data (e.g., compounds may have an associated chemical formula). The respective data models allow for a standardized in-memory integration, storage, and manipulation of data. For example, the reaction data models are especially helpful for integrating the same reaction from multiple databases; since some reaction database entries do not provide cross-linking, it may be necessary to match reaction entities through their stoichiometry and the compound entities they are associated with (i.e., reactants and products). If the stoichiometry and the reactants and products compound entities are the same, the reactions can be considered the same and merged into the same data model, thus avoiding redundancy. Additionally, these entities can be connected to other entities (e.g., a gene can be connected to a protein), and can thus be exported as a network. Entities are connected within the network following the search mode and databases used (Figure 3).

**Figure 3.**
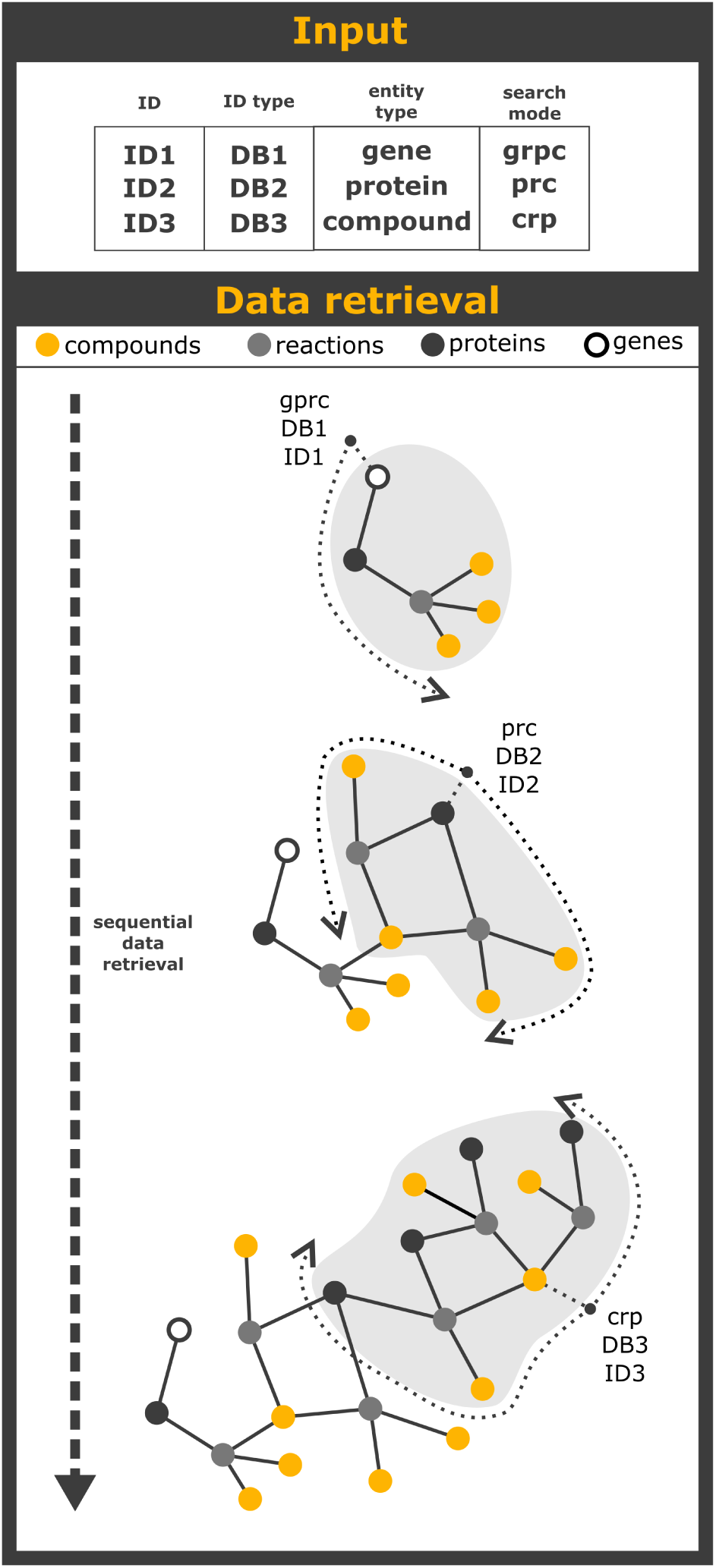
UniFuncNet search modes: Example of three different search modes available in UniFuncNet and how they sequentially link entities, generating a connected network. The first input line contains a gene ID with search mode “gprc”, UniFuncNet searches first for information on this gene and subsequently the directly or indirectly connected entities (one protein, one reaction and three compounds). The second input line contains a protein ID with the search mode “prc”. UniFuncNet retrieves first information on the protein, then on two reactions and four compounds; notice how one of the compounds found in the second search is linked to the network created already during the processing of the first input. The third input line contains a compound ID, and the search mode “crp”, UniFuncNet then retrieves information on four compounds, three reactions and four proteins. Again, since one of the proteins was already searched during the processing of the second input line, the resulting network will connect these inputs’ entities.

#### Search modes

UniFuncNet’s data models represent the four main entity types (”g” = gene, “p” = protein, “r” = reaction, “c” = compound, see above). These data models are then organized to be retrieved according to the underlying structure of each database; i.e., biological databases entities are generally connected in two directions: *g*→*p*→*r*→*c* and *c*→*r*→*p*→*g*.

UniFuncNet can process entities in 14 possible search modes, i.e., “gp”, “gpr”, “gprc”, “pg”, “pr”, “prc”, “rpg”, “rp”, “rc”, “cr”, “crp”, “crpg”, “”, and “global”. Each letter in the search mode corresponds to one of the four different entity types. The “global” search mode corresponds to a search in both directions, e.g., while searching for a given protein, UniFuncNet retrieves information on the associated genes - “pg”, as well as the associated reactions and compounds - “prc”. The “” search mode corresponds to an “*in situ*” search on the same entity, i.e., UniFuncNet retrieves information on the given input IDs without connecting them to additional other entities, e.g., when one aims to fetch ChEBI IDs from compound synonyms or for ID conversion. Figure 3 represents a generic example of multiple search modes and how these drive network generation.

The user input and search mode are inherently linked to the data that is collected, i.e., the user input ID is used as a seed for data retrieval and to generate an entity, whereas the search mode is used to impose a direction and stop criterion on the data retrieval process. If, for example, the user inputs a reaction ID - the resulting entity will contain the database IDs associated with this reaction. During data retrieval this entity may also be connected to different types of entities, e.g., a reaction entity is usually associated with two or more compound entities. The IDs of these connected entities are then used for posterior data retrieval and generation of the respective entities (Figure 3). The user is able to input multiple search modes (comma separated) for the same input ID, which may be useful, e.g., for connecting a reaction entity to its respective compound and protein entities.

#### UniFuncNet to Neo4j API

In order to provide users with the possibility to efficiently query and manage the UniFuncNet results (for example during network analysis), an API importing UniFuncNet’s output into Neo4j a highly-flexible graph database management system, was implemented.

UniFuncNet’s output can be depicted as a multipartite graph, which is a graph whose nodes can be split into multiple independent sets. In the case of UniFuncNet each output file contains multiple entities (e.g., proteins) with entity related annotations (e.g., EC numbers (ECs)) (Figure 4). Since Neo4j is a highly flexible graph-based database it provides a natural integration of UniFuncNet’s data models.

**Figure 4.**
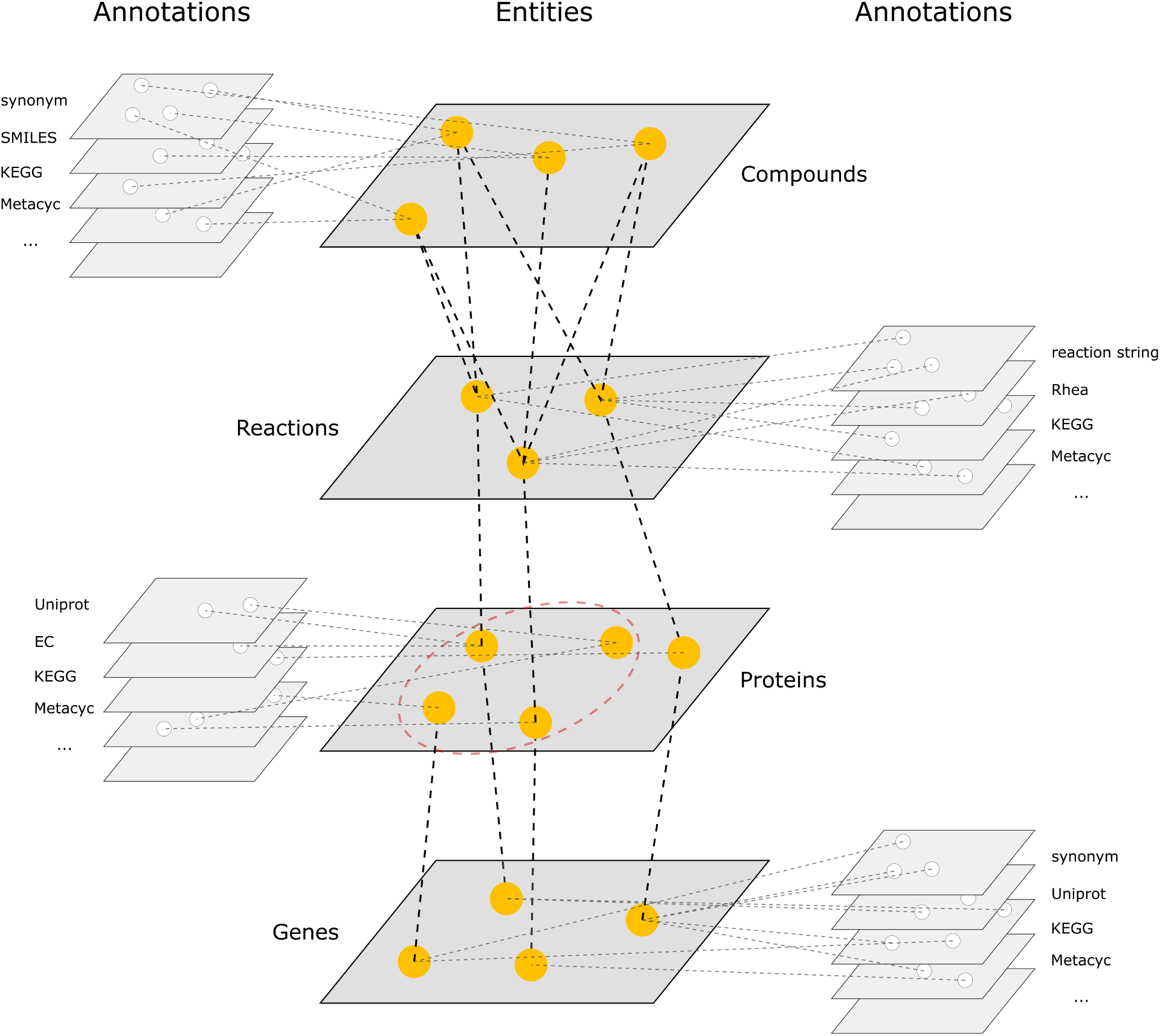
UniFuncNet results as a multipartite graph. The output from UniFuncNet can be represented as a multipartite graph, where the central layers correspond to the entity types (e.g., proteins), and the outer layers to the annotations (e.g., IDs or synonyms). The protein layer contains a protein complex (red dashed circle), comprised of multiple subunits (i.e., protein nodes).

The API takes as input a folder containing all the UniFuncNet output tsv files and stores the data in a Neo4j database. This database can then be queried using Cypher (Neo4j’s querying language) or using any programming language Neo4j API (e.g., the Python or Java drivers). Additionally, we added the option to input Mantis consensus annotations to query the Neo4j database and create the respective SIF networks.

### Use cases

UniFuncNet is a flexible network annotation framework, being usable within diverse contexts. We provide two case scenarios, the first using UniFuncNet for the expansion of GSMMs, and the second for linking compounds with specific organisms.

#### Use case I

In this use case we used UniFuncNet to expand GSMMs built with CarveMe[34], exploring how many putative connections UniFuncNet could add to the original GSMM. To that end, Mantis is used to provide additional functional annotations, and UniFuncNet to map those functional annotations to the GSMM (Figure 1.A).

As an example, we expanded the GSMMs of multiple organisms that have been shown to be relevant in multiple ecosystems; (i) *Akkermansia muciniphila* has been shown to play an important role in human intestinal health as part of the gut microbiome [47]; (ii) *Bradyrhizobium japonicum*, has been shown to be a key organism in nitrogen fixation, essential for e.g. soybean plant growth [48]; (iii) *Microthrix parvicella*, has been shown to be the dominant species involved in the bulking of activated sludge and lipid accumulation in wastewater treatment plants [49]; (iv) *Pelagibacter ubique*, has been shown to be an ubiquitous ocean-dwelling bacterium that belongs to the SAR11 clade, which is reported to account for 25% of all cells in the ocean [38], and (v) *Streptococcus pneumoniae*, has been shown to be a key human pathogen, which is one of the leading causes of pneumonia, bacterial meningitis, and sepsis [50].

This workflow compiled a list of all ECs and MetaCyc protein IDs found by Mantis that are not part of the original GSMM (generated by CarveMe). A non-redundant list of IDs over all species was generated.

This list was converted to a UniFuncNet input file, which contained 1329 unique EC numbers and 1052 unique MetaCyc protein IDs. UniFuncNet were then run for all these IDs with the “prc” search mode to connect the (p)rotein function annotations to the (r)eactions and (c)ompounds.

UniFuncNet collected 5244 putative reactions, which were then filtered according to multiple steps: (i) filter for proteins associated to at least one reaction (ii) filter for proteins that were also absent in the original GSMM (iii) extract all reactions connected to these proteins, (iv) exclude reactions that were already present in the original GSMM, and (v) match the compounds obtained from UniFuncNet with the GSMM compounds to match reactions and exclude redundant reactions.

For each proteome, a baseline directed network from the initial GSMM was created, where reactions and their respective substrates and products are represented as nodes (i.e., reactant(s)→reaction→product(s)). We then expanded the network by adding UniFuncNet’s nodes, either by adding new connections to the baseline network or adding new nodes.

The draft GSMMs and expanded networks were evaluated according to: (i) % of reactions in the largest network component (%RLC); (ii) % of dead end metabolites (%DEM), i.e., metabolites without a transporter reaction that are produced but not consumed or consumed but not produced [51]; (iii) % of newly connected dead end metabolites (%CDEM); and (iv) the amount of new putative reactions that could be added to the GSMM (NR).

On average %RLC decreased from 99.6% (sd=0.4%) to 95.0% (sd=0.7%), &DEMs increased from 4.2% (sd=0.6%) to 12.8% (sd=0.6%), and 0.1% (sd=0.06%) of DEMs were successfully connected in the expanded network. Finally, on average 1005 (sd=485.5) reactions could be added per proteome.

The expanded networks resulted in a substantial enzyme-specific enrichment (i.e., ECs), the most enriched ones being transferases, oxidoreductases and hydrolases. We also analysed the ECs functional redundancy of the each organisms’ baseline and expanded networks, i.e., each baseline network was compared to all others baseline networks, and the same was repeated for the expanded networks. On average, we found that the ECs functional redundancy for the “only baseline”, “only expanded”, “baseline+expanded” networks was 0.74, 0.44, and 0.66 (0-1, 1 being equal), respectively.

When analysing KEGG pathways, the most enriched metabolic capacities corresponded to the metabolism of carbohydrates, lipids, and cofactors and vitamins (from least to most enriched). In the *Akkermansia muciniphila* expanded network, the biosynthesis and metabolism of glycans was among the metabolic capacities most enriched by the network expansion (161 ECs in the baseline to 166 additional ECs in the expanded network mapped to the kegg pathway “Glycan biosynthesis and metabolism”). In the *Microthrix parvicella* expanded network, the metabolism of lipids was the metabolic capacity most enriched by the network expansion (31 ECs in the baseline to 218 additional ECs in the expanded network mapped to the kegg pathway “Lipid metabolism”).

These results are available in Supplementary table “results.ods” available at https://gitlab.lcsb.uni.lu/pedro.queiros/benchmark_unifuncnet.

#### Use case II

In order to understand how UniFuncNet could be used to link different omics levels, we used it to connect functionally annotated reference proteomes to a metabolomics dataset, i.e., linking metabolism related proteins to their reactions and respective compounds (Figure 1.B).

As an example, we used a metabolomics study [41] that cultured four organisms in artificial sputum and nutrient broth mediums and sampled their headspaces for 13 compounds. These compounds were used as potential biomarkers in order to determine the most appropriate antimicrobial therapy in the treatment of ventilator-associated pneumonia.

In order to find the reactions and proteins associated with each compound, UniFuncNet ran with the search mode “crp”. The proteins found to be connected with the compounds via UniFuncNet were then intersected with the functional annotations of each proteome, thus allowing for the identification of the enzymes within each organism involved in the metabolism of these compounds.

After running this workflow we successfully retrieved information on 11 of 13 compounds, eight of these were linked to a total of 30 reactions. These reactions were then connected to a total of 17 proteins. We then linked the proteins connected to reactions (n=17) to the functional annotations of each organism, finding which of these organisms could potentially be involved in the metabolism of studied compounds. We found that all organisms were involved in the metabolism of indole and that *Pseudomonas aeruginosa* was additionally involved with the metabolism of 2-furanmethanol.

## Discussion and conclusion

Here we present UniFuncNet, a network annotation framework that collects and integrates data from multiple biological databases. UniFuncNet can be used to search for information and generate annotated networks in a flexible manner (i.e., various search modes and input ID types). UniFuncNet automates data collection into a human-readable output, by connecting the different biological entities (i.e., genes, proteins, reactions, and compounds), and it provides a network-structured output, which can be easily used in network-based downstream analysis. An added benefit of UniFuncNet is the standardization of the search methodology, potentially decreasing the accidental omission of information during manual collection/curation.

UniFuncNet collects data from live websites/application programming interfaces (API) and allows the user to update their own local flat files (e.g., MetaCyc or Rhea). UniFuncNet ensures that the collected data is up to date, which represents a limitation in similar projects [46]), since they require regular database maintenance. However, UniFuncNet faces its own challenges: (i) a website’s HTML structure or API may change over time, which requires maintenance of UniFuncNet’s data collection protocols, (ii) live retrieval of information tends to be slower than using a centralized source of data, and (iii) websites may block scraping attempts if these are done too frequently, which is circumvented by UniFuncNet by having 10 second waiting period between each web query to the same database. While reliable and large data collection is provided by UniFuncNet, as a framework that can speed-up the work of researchers requiring comprehensively annotated networks, it is advisable to perform manual curation during downstream data integration. Overall though, we believe that the benefits of having a lightweight framework with very low storage footprint, always retrieving the latest information, clearly outweigh the aforementioned downsides.

Current automated reconstruction tools are capable of generating GSMMs ready for modelling. However, divergent implementations [52, 34, 53] may lead to different outcomes (i.e., the models) due to multiple factors, e.g.: (i) different gene predictions, (ii) different functional annotation reference databases, and (iii) different automated curation implementations [54, 55]. While automated curation offers a good modelling basis, it is unlikely that the current methods will ever be able to encompass the complexity of *in vivo* biological networks. As such, manual curation and expansion of GSMMs remain essential; the latter is routinely done through the iterative analysis of the subsystems for genes, proteins, and reaction(s) of interest. In particular, the end-user searches for information regarding a certain ontology ID, such as KEGG [18] orthology IDs, ECs, or others, in highly comprehensive (and partially redundant) biological databases. To avoid introducing redundancy, cross-linking entities between databases is necessary, which can be done manually or partially automated through ID mapping tools offered by MetaNetX [46] or UniProt [27]. To this end, we implemented a workflow that uses UniFuncNet to facilitate the cross-linking and expansion of GSMMs.

We have shown that the networks enriched with UniFuncNet’s workflow were better able to capture organism-specific characteristics, e.g., in *Microthrix parvicella* the metabolism of lipids was the most enriched KEGG pathway, which agrees with the findings of Sheik et al. [49]. Similarly, in *Akkermansia muciniphila* the metabolism and biosynthesis of glycans was amongst the most enriched KEGG pathways, which supports the hypothesis that this organism and glycans play an important role in human gut health [47, 56]. Lastly, the metabolism of cofactors and vitamins was, on average, the most enriched KEGG pathway among all organisms.

In general, we found that this workflow could add a substantial amount of reactions to the GSMMs, which, as previously shown [31], is likely due to the more comprehensive reference databases used (Mantis with the KOfam, Pfam, and MetaCyc databases and CarveMe with the BIGG database [57]). In addition, we also found that the similarity between the functional profiles (i.e., ECs functional redundancy) between each organism’s network was substantially lower in the expanded networks, highlighting the benefit of applying UniFuncNet to discover functions unique to each organism. It is important to emphasize that the expanded networks would still require curation; indeed the aim of this workflow is not to directly output an expanded GSMM ready for modelling, but to provide the user with a framework that automates some of the most time-consuming curation steps, i.e., expanding and enriching the network. Altogether, these results show the potential of UniFuncNet to support the expansion of GSMMs, provided additional curation steps are implemented by the end-users. While in this manuscript we have shown how UniFuncNet can be used in a targeted manner, it could also be used for the generation of genome-scale metabolic networks.

We have also shown how UniFuncNet can be used to link different datasets, in particular how it can be used for linking different omics, which should prove useful for multi-omics network-based analysis. Specifically, in the second workflow, we have shown how UniFuncNet may be used for the mapping of compounds to specific organisms. The results shown in the use case II were not able to connect the organisms and compounds in the same resolution as the respective study [41], which further highlights the need to create and use more comprehensive functional annotation reference databases. However, we found that indole’s metabolism was shared among all organisms, which is a clear indication of conservation of function in prokaryotes[58, 59, 60]. Despite this, we believe this workflow could be combined with more resolved input proteomes (i.e., using proteomics data instead of reference proteomes) and as such could be an even more powerful screening tool for more thorough investigations.

In conclusion, in this article we have highlighted UniFuncNet’s ability to automatically and comprehensively annotate networks. Additionally, we have showcased two use cases which could be used as baseline examples for more intricate analysis. We believe that UniFuncNet’s flexible search modes and varied input formats expands its utility into a variety of analysis well beyond the ones shown in this paper.

## Acknowledgements

All authors proof-read and approved of the content in this research paper. The authors declare that they have no competing interests. We would like to acknowledge Ines Thiele and Alberto Noronha for their supervision during the initial stages of this project. Supported by the Luxembourg National Research Fund PRIDE17/11823097 and the European Research Council (ERC) under the European Union’s Horizon 2020 research and innovation programme (grant agreement No. 863664).

## Conflict of interest statement

None declared.

## Notes

https://github.com/PedroMTQ/UniFuncNet

https://gitlab.lcsb.uni.lu/pedro.queiros/benchmark_unifuncnet

### Competing Interest Statement

The authors have declared no competing interest.

### Summary of Updates

Author order updated in the biorxiv's author list added figures

